# ERAP1 allotypes 2 and 10 differentially regulate the immunopeptidome of melanocytes

**DOI:** 10.1101/2025.05.08.652898

**Authors:** Martha Nikopaschou, Martina Samiotaki, Anna Kanavou, Nikos Angelis, Ourania Tsitsilonis, George Panayotou, Efstratios Stratikos

**Affiliations:** National Centre for Scientific Research Demokritos, Agia Paraskevi, Greece; Department of Chemistry, National and Kapodistrian University of Athens, 15784 Zografou, Greece; Biomedical Sciences Research Center “Alexander Fleming”, Institute for Bioinnovation, 16672 Vari, Greece; Department of Biology, National and Kapodistrian University of Athens, 15784 Zografou, Greece

## Abstract

Endoplasmic reticulum aminopeptidase 1 (ERAP1) is a polymorphic enzyme that shapes the peptide repertoire presented by MHC class I molecules and can regulate adaptive immune responses in cancer and autoimmunity. Common missense polymorphisms in ERAP1 modulate its activity and are found in specific allotypes in humans. ERAP1 allotypes are linked to predisposition to HLA-associated inflammatory diseases. ERAP1 allotypic variation has been correlated with the development of psoriasis and Bechet’s disease, through the generation of specific CD8+ T cell populations targeting disease-specific HLAs. Given the established broad effects of ERAP1 activity on the cellular immunopeptidome, we hypothesized that ERAP1 allotypic variation may lead to broad immunopeptidome shifts that underlie antigenic responses. To test this hypothesis, we generated two A375 melanoma cell lines, each one expressing one of the most common, disease-associated, ERAP1 allotypes, namely allotypes 2 or 10. Comparison of the immunopeptidome of these two cell lines showed only minor differences in peptide sequences presented but extensive changes in abundance that included alterations in length distribution, binding affinity and sequence motifs. Our results suggest that enzymatic differences between ERAP1 allotypes are reflected primarily on the quantitative composition of the cellular immunopeptidome. These quantitative changes may constitute a mechanism that underlies ERAP1-allotypic associations with HLA-associated autoimmunity and variable anti-tumor responses.

## INTRODUCTION

Cellular adaptive immune responses are primarily mediated by specialized immune cell receptors that recognize Major Histocompatibility Class I molecules (MHC-I; HLA in humans) complexed with small peptides derived from the proteolytic degradation of intracellular proteins^1^. These peptides are generated by complex intracellular proteolytic cascades and can encompass thousands of different amino acid sequences, collectively known as the immunopeptidome^2^.

Peptides originating from self proteins do not normally elicit immune responses. This happens primarily due to the clonal deletion of T cell clones carrying receptors that can recognize them during the development of immunological tolerance^3^. On the contrary, peptides originating from infection or oncogenic transformation of the cell can be recognized by immune cells, such as CD8+ T cells, leading to the eradication of the antigen presenting cell^4^. Indeed, presentation of cancer-associated antigens is a major determinant of anti-tumor immunity and often a target in immunotherapy efforts^5,6^.

Insufficient formation of tolerance, chronic inflammation and pathogen antigenic mimicry can sometimes lead to the erroneous recognition of self-peptides as pathogenic, initiating and/or sustaining autoimmune inflammatory responses and leading to disease^7–9^. Such a molecular mechanism has been suggested to underlie the pathogenesis of a group of diseases termed MHC-I-opathies, such as Ankylosing Spondylitis, Spondylarthritis, Behçet’s disease, Birdshot Uveitis and psoriasis^10^. The HLA in humans is highly polymorphic with more than 20000 different haplotypes described^11^. MHC-I-opathies have a very strong genetic linkage with specific HLA alleles, suggesting possible pathogenic mechanisms that involve specific antigenic peptides or sets of antigenic peptides presented by the disease-associated HLA^12^. Indeed, recent work has identified antigenic peptide recognition in the context of disease-associated HLA by specific T cell receptors on CD8+ cells, providing further support for this pathogenic hypothesis^13–15^.

A key regulatory node shaping the cellular immunopeptidome is an endoplasmic reticulum (ER)-resident enzyme called ER aminopeptidase 1 (ERAP1)^16^. ERAP1 trims N-terminal amino acids from extended precursors of mature antigenic peptides, bringing them to optimized lengths for binding to nascent MHC-I^17^. Although ERAP1 preferentially trims larger peptides, it can often over trim mature antigenic epitopes so that they are no longer suitable for MHC-I binding, essentially destroying them^18^. By this dual function, the activity of ERAP1 can regulate the immunopeptidome and adaptive immune responses^19–21^. Thus, ERAP1 is currently a pharmacological target aiming to regulate the antigenicity of cells, in applications such as cancer immunotherapy^22,23^.

Like HLA, ERAP1 is also polymorphic and several single nucleotide polymorphisms (SNPs) have been associated with predisposition to human disease, including viral infections, anti-tumor responses and most notably MHC-I-opathies^24–26^. Often, genetic predisposition between ERAP1 SNPs is found to be in epistasis with HLA alleles, consistent with the molecular role of the enzyme in generating the peptide repertoire that is available to bind to MHC-I^27^. While some ERAP1 SNPs affect protein expression^28^, many result in missense mutations generating ERAP1 variants with distinct enzymatic properties^29^. Within the human population, these coding SNPs organize in specific allotypes that present specific functional properties ^30,31^. Of note, ERAP1 allotype 10 was first reported to be a loss-of-function variant and exists in about a quarter of the European population^32^. Follow-up studies, however, have suggested that ERAP1 allotype 10 rather presents a unique specificity for peptide substrates^31,33^.

A recent report identified a T cell receptor from psoriasis patients’ T cells recognizing an autoantigen presented by HLA-C*06:02 in melanocytes^13^. This autoantigen required ERAP1 for its generation and this process was dependent on ERAP1 allotypic variation. Allotype 2 was highly efficient in generating the autoantigenic peptide and inducing a T cell response, while allotype 10 was ineffective and reduced immunogenicity. This finding was consistent with genetic predisposition findings which suggested that ERAP1 allotype 2 is predisposing to psoriasis while allotype 10 is protective and introduced a molecular framework for understanding these effects. In addition, ERAP1 allotype 10 has been shown to change the immunogenicity of antigenic peptides presented by the Bechet’s disease associated HLA-B*51 allele^34,35^. Since the enzymatic activity of ERAP1 has been demonstrated to also affect the immunogenicity of cancer cells and responses to immunotherapy, the differential activity of allotypes may also regulate anti-tumor responses^25,36–38^.

Given the complex landscape of allotype 10 functional differences, we hypothesized that changes in antigen presentation between ERAP1 allotypes may not be limited to single antigens but rather extend to broad shifts in the immunopeptidome, which can have implications for both autoimmunity and cancer. To test this hypothesis, we generated monoallelic clones of the melanoma cell line A375 that carry either ERAP1 allotype 2 or allotype 10; these are the most common allotype world-wide and the most common allotype in European populations^30^, respectively, and have been associated with autoimmunity^13,34,35^.

Analysis of the immunopeptidomes of these two clones revealed broad differences in the abundance of presented peptides that comprise changes in length distribution, sequence motifs and binding affinity, all of which can influence recognition by immune cell receptors. These results suggest that functional enzymatic differences between ERAP1 allotypes are reflected in the cellular immunopeptidome and may underlie mechanisms linking ERAP1-mediated antigen processing to HLA-associated autoimmunity and cancer.

## EXPERIMENTAL PROCEDURES

### Plasmids

pcDNA3.1(+) vectors carrying the codon-optimized coding sequence for ERAP1 allotype 2 and allotype 10 were obtained by custom gene synthesis (BioCat GmbH, Heidelberg). The ERAP1 gene was inserted in pcDNA3.1(+) between BamHI and XhoI cloning sites. The amino acid sequences for both allotypes are shown below (signal sequence is underlined, and differences in SNPs are indicated by bold).

#### Allotype 2

<u>MVFLPLKWSLATMSFLLSSLLALLTVSTPSWC</u>QSTEASPKRSDGTPFPWNKIRLPEYVIPVH YDLLIHANLTTLTFWGTTKVEITASQPTSTIILHSHHLQISRATLRKGAGERLSEEPLQVLE HPRQEQIALLAPEPLLVGLPYTVVIHYAGNLSETFHGFYKSTYRTKEGELRILASTQFEPTA ARMAFPCFDEPAFKASFSIKIRREPRHLAISNMPLVKSVTVAEGLIEDHFDVTVKMSTYLVA FIISDFESVSKITKSGVKVSVYAVPDKINQADYALDAAVTLLEFYEDYFSIPYPLPKQDLAA IPDFQSGAMENWGLTTYRESALLFDAEKSSASSKLGIT**M**TVAHELAHQWFGNLVTMEWWNDL WLNEGFAKFMEFVSVSVTHPELKVGDYFFGKCFDAMEVDALNSSHPVSTPVENPAQIREMFD DVSYDKGACILNMLREYLSADAFKSGIVQYLQKHSYKNTKNEDLWDSMASICPTDGVKGMDG FCSRSQHSSSSSHWHQEGVDVKTMMNTWTLQ**K**GFPLITITVRGRNVHMKQEHYMKGSDGAPD TGYLWHVPLTFITSKS**D**MVHRFLLKTKTDVLILPEEVEWIKFNVGMNGYYIVHYEDDGWDSL TGLLKGTHTAVSSNDRASLINNAFQLVSIGKLSIEKALDLSLYLKHETEIMPVFQGLNELIP MYKLMEKRDMNEVETQFKAFLIRLLRDLIDKQTWTDEGSVSE**R**MLRS**Q**LLLLACVHNYQPCV QRAEGYFRKWKESNGNLSLPVDVTLAVFAVGAQSTEGWDFLYSKYQFSLSSTEKSQIEFALCRTQNKEKLQWLLDESFKGDKIKTQEFPQILTLIGRNPVGYPLAWQFLRKNWNKLVQKFELGSSSIAHMVMGTTNQFSTRTRLEEVKGFFSSLKENGSQLRCVQQTIETIEENIGWMDKNFDKIRVWLQSEKLERM

#### Allotype 10

<u>MVFLPLKWSLATMSFLLSSLLALLTVSTPSWC</u>QSTEASPKRSDGTPFPWNKIRLPEYVIPVH YDLLIHANLTTLTFWGTTKVEITASQPTSTIILHSHHLQISRATLRKGAGERLSEEPLQVLE HPPQEQIALLAPEPLLVGLPYTVVIHYAGNLSETFHGFYKSTYRTKEGELRILASTQFEPTA ARMAFPCFDEPAFKASFSIKIRREPRHLAISNMPLVKSVTVAEGLIEDHFDVTVKMSTYLVA FIISDFESVSKITKSGVKVSVYAVPDKINQADYALDAAVTLLEFYEDYFSIPYPLPKQDLAA IPDFQSGAMENWGLTTYRESALLFDAEKSSASSKLGIT**V**TVAHELAHQWFGNLVTMEWWNDL WLNEGFAKFMEFVSVSVTHPELKVGDYFFGKCFDAMEVDALNSSHPVSTPVENPAQIREMFD DVSYDKGACILNMLREYLSADAFKSGIVQYLQKHSYKNTKNEDLWDSMASICPTDGVKGMDG FCSRSQHSSSSSHWHQEGVDVKTMMNTWTLQ**R**GFPLITITVRGRNVHMKQEHYMKGSDGAPD TGYLWHVPLTFITSKS**N**MVHRFLLKTKTDVLILPEEVEWIKFNVGMNGYYIVHYEDDGWDSL TGLLKGTHTAVSSNDRASLINNAFQLVSIGKLSIEKALDLSLYLKHETEIMPVFQGLNELIP MYKLMEKRDMNEVETQFKAFLIRLLRDLIDKQTWTDEGSVSE**Q**MLRS**E**LLLLACVHNYQPCV QRAEGYFRKWKESNGNLSLPVDVTLAVFAVGAQSTEGWDFLYSKYQFSLSSTEKSQIEFALCRTQNKEKLQWLLDESFKGDKIKTQEFPQILTLIGRNPVGYPLAWQFLRKNWNKLVQKFELGSSSIAHMVMGTTNQFSTRTRLEEVKGFFSSLKENGSQLRCVQQTIETIEENIGWMDKNFDKIRVWLQSEKLERM

### Generation of Cell lines

A375 cells ERAP1 Knock-Out (KO) cells (clone 1B12, previously described^38^) were cultured in high glucose DMEM (Biowest, L0104) with the addition of 10% Fetal Bovine Serum (Biowest, S1810), 2 mM L-glutamine (Biowest, X0550) and 1% Penicillin-Streptomycin (Biowest, L022) as usual at 37 °C, 5% CO_2_. For the generation of A375 clones, the pcDNA3.1(+) plasmids were linearized with SspI-HF® (New England Biolabs, R3132) and purified with Monarch® PCR & DNA Cleanup Kit (New England Biolabs, T1030), prior to transfection of A375 ERAP1 KO cells. For the transfection, A375 KO cells were seeded in a 6-well plate (500,000 cells/well) and 24 hrs later they were transfected with each vector, using the jetPRIME® (Polyplus, 101000027) transfection reagent, according to the manufacturer instructions. 48 hours post-transfection, both transfected and untransfected cells were exposed to 1.2 mg/ml G-418 (InvivoGen, PHC4033) (established from an antibiotic killing curve on untransfected A375 KO cells) for 2 weeks, with frequent medium changes. Surviving cells were diluted in 96-well plates (aiming for 0-1 cells/well) to obtain single clones, which were then allowed to grow. Western blots using a human aminopeptidase PILS/ARTS1 antibody (R&D Systems, AF2334) were performed to confirm the ERAP1 expressing clones and to identify clones with similar levels of expression for the two allotypes. Positive clones were cultured in the complete medium stated above but also supplemented with 0.6 mg/ml G-418.

### Flow cytometry

Selected positive clones (2B1 and 1H1) were seeded in a 24-well plate (100,000 cells/well) and 24hrs later they were treated with human recombinant interferon-γ (IFN-γ, Gibco, PHC4033) at concentrations ranging from 0 to 50 ng/ml for another 24 hrs. Cells were subsequently detached and prepared for flow cytometry (FC) analysis, after staining with anti-human HLA-ABC FITC labelled antibody (Biorad, MCA81F) (1:25). FACS analysis was performed with a FACSCelesta™ cell analyzer (BD Biosciences) using the FACSDiva™ Software (BDBiosciences).

### Isolation of the immunopeptidome

The immunopeptidome of A375 cells was isolated as previously described^39^. Cells were grown in three separate biological replicates, treated with 20 ng/ml of IFN-γ for 24 hrs, detached using Accutase (Merck Millipore, SCR005), pelleted and stored at -80°C (2.5×10^8^ per sample). For isolation of MHC-I, cell pellets were thawed on ice and cells lysed as described^39^ with some modifications. Briefly, samples were lysed with 10 ml lysis buffer/sample (Tris–HCl, pH 7.5, 150 mM NaCl, 0.5% IGEPAL CA-630, 0.25% sodium deoxycholate, 1 mM EDTA, pH 8.0, cOmplete™ EDTA-free protease inhibitor cocktail tablets). The lysate was then cleared with ultracentrifugation at 100000 g for 1 hour at 4 °C, prior to loading on CN-Br pre-columns followed by the W6/32 coupled columns (generated by coupling W6/32 antibody, grown in-house, onto cyanogen bromide activated Sepharose 4B beads, as previously described^39^). The flow-through from this procedure was loaded on the columns three more times prior to washing of the columns with 20 bed volumes 20 mM Tris–HCl, pH 8.0, 150 mM NaCl, 20 bed volumes 20 mM Tris–HCl, pH 8.0, 400 mM NaCl, 20 bed volumes 20 mM Tris–HCl, pH 8.0, 150 mM NaCl and finally with 40 bed volumes 20 mM Tris–HCl, pH 8.0. MHC-I-peptide complexes were eluted by washing with 1% trifluoroacetic acid (TFA) and the samples were stored at -80 °C. Peptide eluates were subjected to SpeedVac and separated from MHC-I molecules using reversed-phase C18 disposable spin columns (Thermo Scientific, 89870). Adequate separation was evaluated with a western blot against β-2 microglobulin (R&D Systems, MAB8248). Ultimately, the purified peptides were further processed by the Sp3 protocol for peptide clean-up, as described previously^39^, solubilized in the mobile phase A (0.1% FA in water) and sonicated.

### Liquid chromatography/ mass spectrometry

For LC-MS/MS analysis, a setup consisting of a Dionex Ultimate 3000 nano RSLC online with a Thermo Q Exactive HF-X Orbitrap mass spectrometer was used and data were obtained both in a data-dependent (DDA, 1 technical replicate/sample) and data-independent acquisition (DIA, 2 technical replicates/sample) modes. Samples were initially injected in a 25 cm-long analytical C18 column (PepSep, 1.9μm^3^ beads, 75 μm ID) with a 30-minute gradient. At the start of the gradient, the mobile phase was composed of 7% Buffer B (0.1% FA in 80% ACN). The gradient was then increased to 35% B over 16.5 minutes, followed by a rise to 45% within 1.5 minutes, and then to 99% B over 0.5 minutes. Flow was held stable at 99% B for 1 minute before returning to 7% over the next 10.5 minutes.

For DDA data, a range between 275 and 1100 m/z was scanned in the full MS, with 120K resolving power, AGC of 3*10^6^ and max IT of 100 ms. For the MS/MS, the 5 most abundant ions were selected, resolving power was set to 30K, AGC to 1*10^6^, max IT to 50 ms and NCE to 30.

For the DIA data, a range between 375 and 1100 m/z was selected in the full MS with 120K resolving power, AGC of 3*10^6^ and max IT of 60 ms. For the MS/MS 8 Th windows (39 loop counts) were selected, resolving power was set to 15K, AGC to 3*10^6^, max IT to 22 ms and normalized collision energy to 26.

### Data analysis

MS/MS spectra were searched against the UniProt database (HUMAN_UP000005640_9606, 20,597 entries, retrieved: 24.11.2023) with the non-specific HLA-DIA workflow in Fragpipe ^40^ v 22. In this workflow, which is specifically designed for immunopeptidomics, a hybrid spectral library was produced by DDA and DIA runs and used for a non-specific search with MSFragger for peptides between 7 and 25 amino acids, accounting for cysteinylation as a variable modification. PSMs, ions and peptides were filtered to 1% FDR at each level. Quantification was ultimately performed from the DIA files with DIA-NN^41^. Data were analyzed and visualized with FragPipe Analyst^42^, without further normalization and imputation. Peptides with log2 difference ≥1 and q-value ≤0.05 were considered differentially expressed. MHCMotifDecon 1.0^43^ was used for motif extraction and binding affinity predictions via NetMHCPan 4.1^44^.

## RESULTS

### Generation of the cellular model system

We have previously shown that the A375 melanoma cell line is a good model system for studying the effects of ERAP1 on the immunopeptidome^19,38,39^. Here we used A375 ERAP1 KO cells to generate mono-allelic ERAP1 clones. To achieve this, A375 ERAP1 KO clone 1B12^39^ was transfected with linearized pcDNA3.1(+) vectors carrying either ERAP1 allotype 2 or allotype 10 and treated with G418 to eliminate untransfected cells. Single clones were isolated and expanded and western blots were used to evaluate ERAP1 expression levels (Supplemental Figure 1). Since ERAP1 expression levels can affect the enzymatic activity in the cell, we selected clones 2B1 for allotype 2 and 1H1 for allotype 10 for further experiments, since they had comparable ERAP1 expression levels.

**Figure 1:**
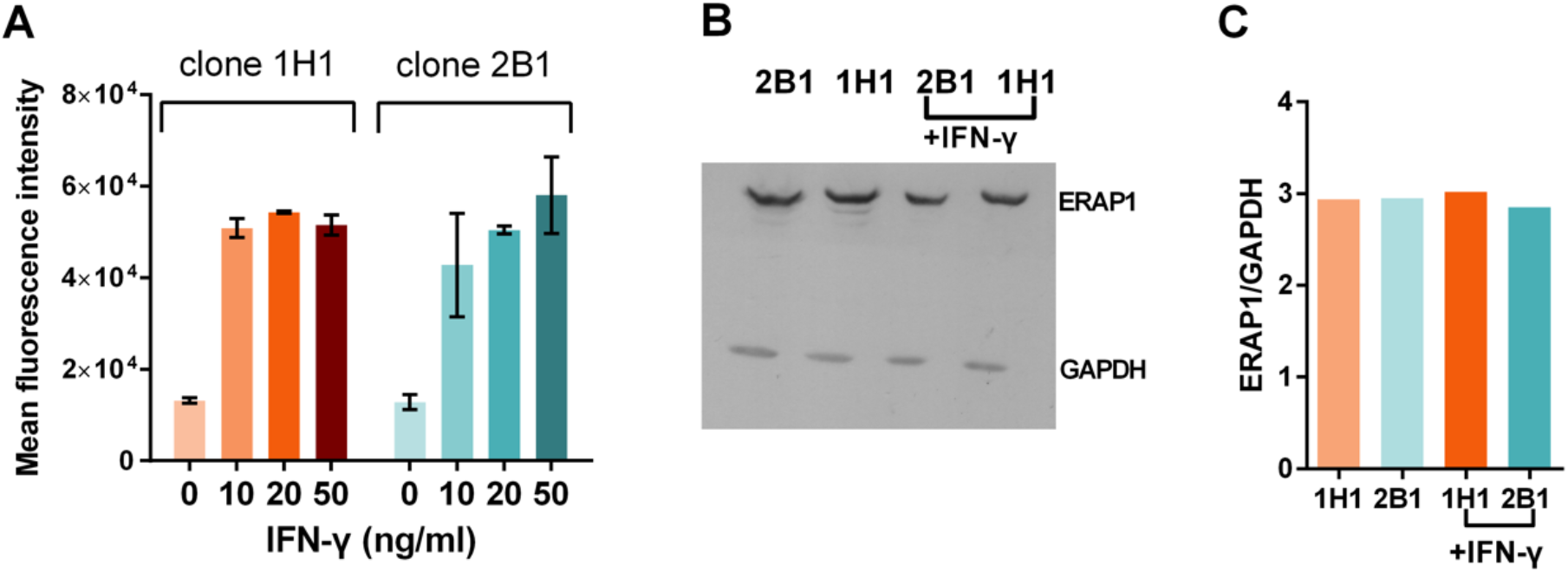
Effects of IFN-γ on cell-surface HLA expression and ERAP1 expression. **Panel A**, Flow cytometry analysis of A375 single ERAP1 allotype clones showing cell-surface HLA levels detected by the W6/32 antibody after incubation with increasing amounts of IFN-γ. Shown are means ± SDs from 3 experiments performed. **Panel B**, western blot analysis of ERAP1 from whole cell lysates in the presence and absence of IFN-γ. **Panel C**, quantification of ERAP1 expression from data shown in panel B, normalized for GAPDH.

Antigen presentation in cancer and autoimmunity often operates in the context of inflammation^13,45^ and cytokines such as interferon-gamma (IFN-γ), can upregulate several components of the antigen presentation machinery, including HLA and ERAP1^46^, enhancing adaptive immune responses. We therefore decided to compare the immunopeptidome of the two A375 clones under the effect of IFN-γ. To optimize the concentration of IFN-γ, we treated the cells with 0, 10, 20 and 50 ng/ml and used FACS to monitor changes in MHC-I cell surface expression (Figure 1A). IFN-γ treatment led to the upregulation of cell surface MHC-I that plateaued at concentrations over 20 ng/ml. As a result, we selected this concentration for immunopeptidome analysis. Cells were also tested for any changes in ERAP1 expression after IFN-γ treatment, since differential upregulation of ERAP1 allotypes would bias the immunopeptidome analysis results. Western blot analysis indicated that ERAP1 expression was not affected by IFN-γ treatment, as expected, given that the integration of the ERAP1 gene after transfection is unlikely to occur in genomic regions under IFN-γ control (Figure 1B,C).

### Isolation and analysis of the immunopeptidome of A375 cells

To analyze the immunopeptidome of the two A375 clones, cells were grown in three separate biological replicates and were treated with 20 ng/ml of IFN-γ for 24 hrs. MHC-I-peptide complexes were then isolated by affinity chromatography from 2.5×10^8^ cells using the W6/32 antibody, as previously described^39^. Eluted peptides from MHC-I complexes were sequenced by LC-MS/MS using both data-dependent and data-independent acquisition. Two technical replicates in data-independent acquisition were performed per biological replicate, totaling 6 replicates for each clone. Obtained MS spectra were then searched against the UniProt database (HUMAN_UP000005640_9606, 20,597 entries, retrieved: 24.11.2023) using the Fragpipe platform^40^. Searching the obtained DIA spectra with the hybrid spectral library built with both DDA and DIA runs resulted in the identification of 2408 peptides between 7 and 25 amino acids (Supplemental Table 1). Principle component analysis indicated that the immunopeptidomes of the two clones formed distinct clusters with wide separation across principal component 1 (Figure 2A). Hierarchical clustering analysis confirmed this observation, as the c distinct clusters (Figure 2B). Thus, this preliminary analysis suggested that the two ERAP1 allotypes produced distinct immunopeptidomes.

**Figure 2:**
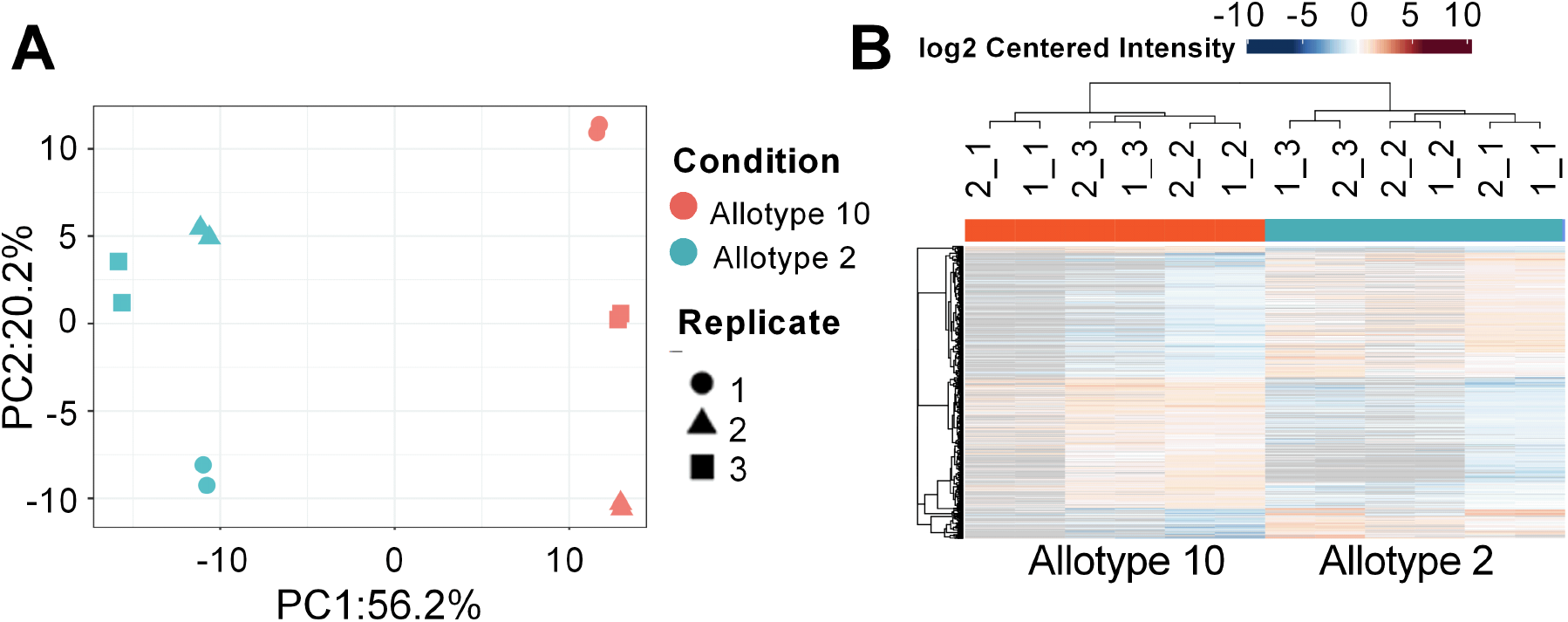
Immunopeptidome analysis of A375 cells. **Panel A**, principal component analysis of the immunopeptidome isolated from the two clones carrying ERAP1 allotypes 2 and 10 (3 biological replicates, 2 technical replicates each). **Panel B**, heatmap of cluster analysis of identified peptides for both experimental conditions and replicates. Color scale indicates peptide intensities (red = high, blue = low).

### The two ERAP1 allotypes induce minor qualitative but significant quantitative immunopeptidome shifts

Comparison of the identified peptide sequences indicated that the majority of peptides were shared between the two immunopeptidomes: 2218 peptides were common in both cell clones carrying the two ERAP1 allotypes, 154 were unique to allotype 2 and 36 to allotype 10 (Figure 3A). This finding was in stark contrast to previous analysis with the same cell line using ERAP1 KO or ERAP1 inhibitors that had revealed major changes in the sequences of detected peptides^19,38^. Statistical analysis, however, revealed significant changes in the abundance of peptides identified in the two conditions (Figure 3B). Specifically, 459 peptides were found to be upregulated in allotype 2 and 408 peptides were upregulated in allotype 10. Combining the differentially upregulated with the unique peptides for each allotype, we calculated that of the total 2408 peptides identified, 613 peptides (25.5 %) were upregulated in allotype 2 and 444 peptides (18.4 %) were upregulated in allotype 10 (Figure 3C). This finding suggests a substantial immunopeptidome shift between cells carrying distinct ERAP1 allotypes.

**Figure 3:**
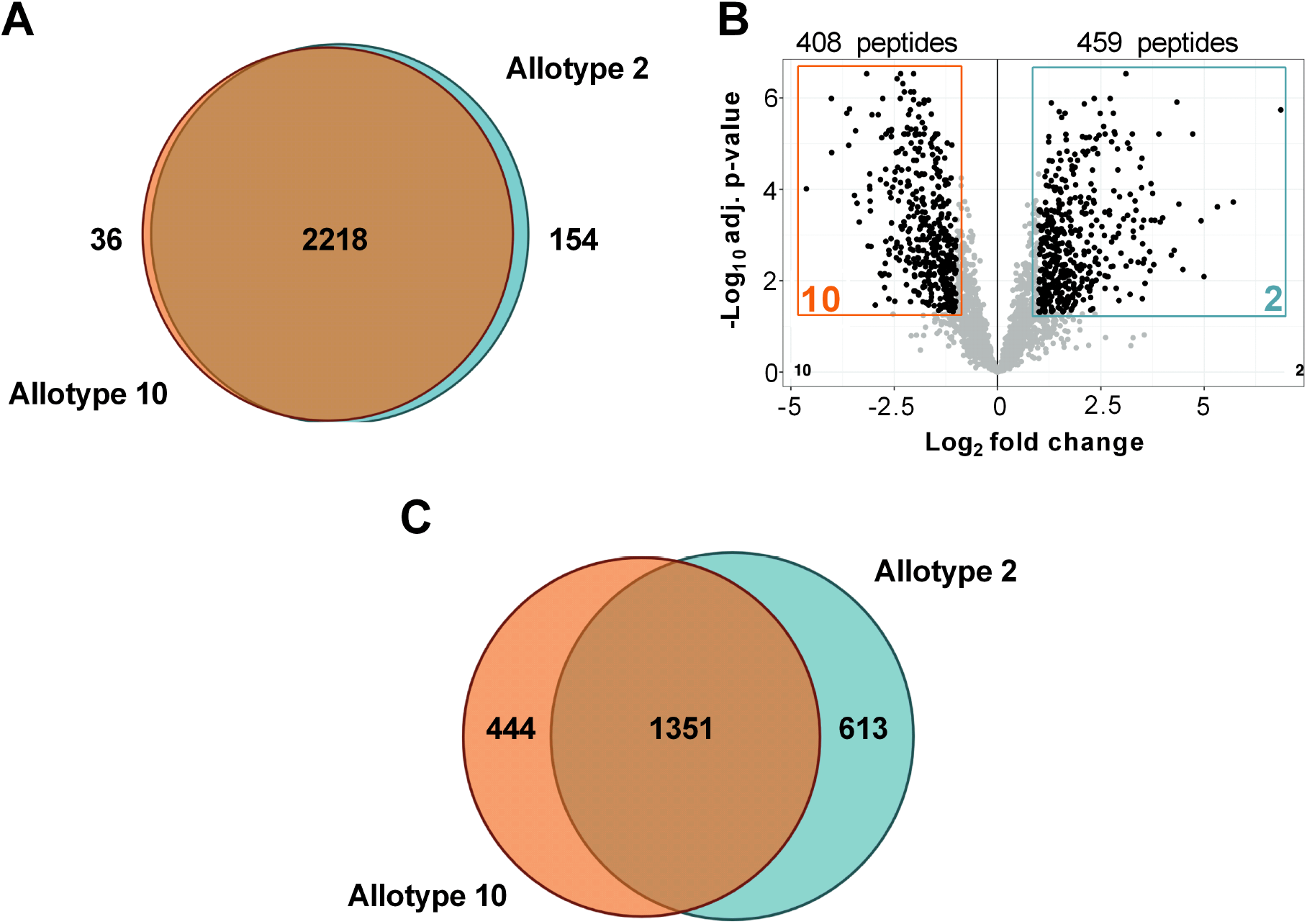
Quantification of immunopeptidome shifts between cells carrying ERAP1 allotype 2 or 10. **Panel A**, Venn diagram depicting the overlap of identified peptide sequences between the two experimental conditions. **Panel B**, Volcano plot showing differences in abundance for identified peptides related to the statistical significance of the observation. Each dot corresponds to a unique peptide sequence. Peptide groups that show statistically significant differences between allotypes 2 and 10 are highlighted with colored boxes. **Panel C**, Venn diagram depicting the immunopeptidome shift between cells carrying ERAP1 allotype 2 or 10, including both uniquely detected peptides and statistically significant abundance changes.

### Differences in affinity, length and HLA allele utilization

To better understand the differences between peptides presented in each of the two immunopeptidomes, we used MHCMotifDecon 1.0^47^ to analyze all the identified peptides between 8 and 14 amino acids. 94.5% of the peptides were predicted to bind to one of the HLA alleles expressed in A375 cells (rank ≤ 2, NetMHCPan 4.1^44^) and corresponded to expected motifs for the respective HLA (Figure 4A,B). Surprisingly, we identified only a small number of peptides with motifs suitable for binding to HLA-C*06:02, a major predisposition allele for psoriasis, and no peptides from the ADAMTSL5 autoantigen previously linked with CD8+ responses in other melanocyte cell lines^13^. The average predicted affinity was found to be better for the peptides presented in cells carrying ERAP1 allotype 10 (Figure 5A). Given that allotype 10 has been reported to have lower enzymatic activity, this finding may underscore the epitope-destructive role of ERAP1 for the HLA alleles carried by this cell line.

**Figure 4:**
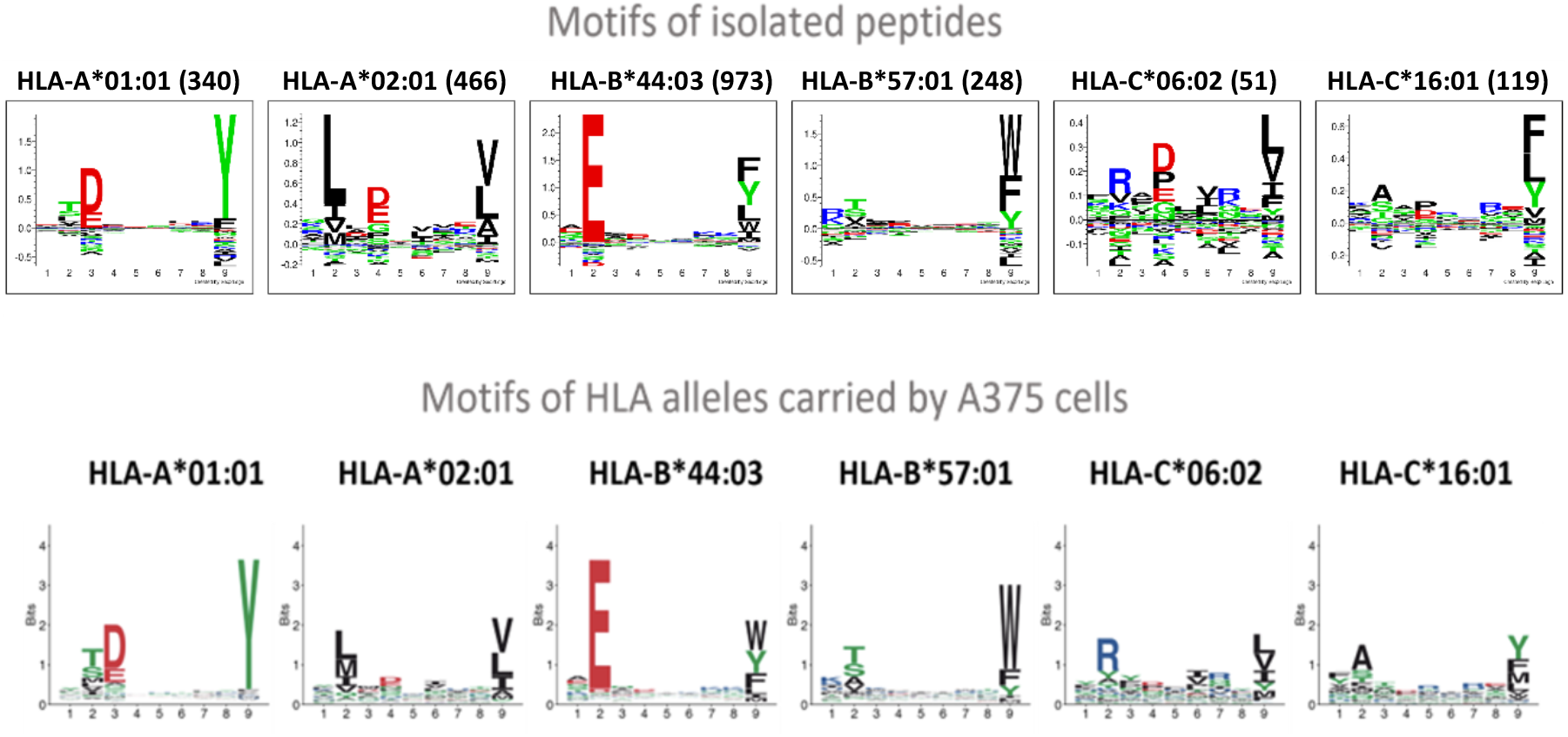
Motifs of identified peptides. **Top**, sequence motifs of identified peptides assigned to one of the HLA alleles carried by A375 cells. **Bottom**, expected sequence motifs for each HLA allele present in A375 cells.

**Figure 5:**
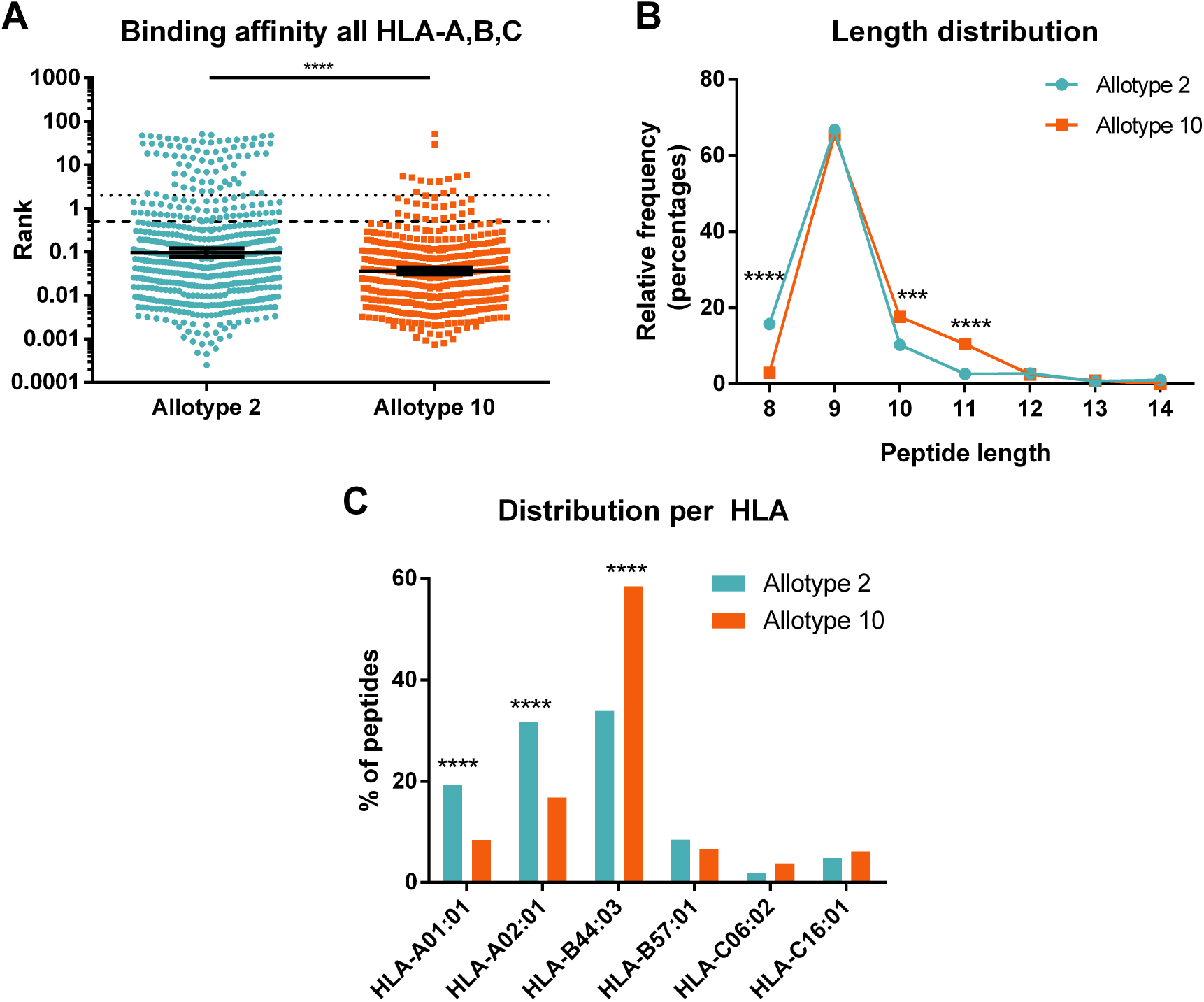
Characteristics of identified peptides from cell lines expressing different ERAP1 allotypes. **Panel A**, predicted binding affinity for at least one of the HLA alleles carried by A375 cells. Each dot corresponds to a different peptide sequence. Peptides below the dotted line are considered binders. The geometric mean of predicted rank is indicated, along with the 95% confidence interval. Statistical significance was evaluated with a Mann-Whitney test in GraphPad Prism v8. **Panel B**, length distribution of identified peptides for the two allotypes. Statistical significance was evaluated with a Chi-square test with Bonferroni correction for multiple testing. **Panel C**, binding assignments to HLA alleles carried by A375 cells. Statistical significance was evaluated with a Chi-square test with Bonferroni correction for multiple testing. p <0.05 (*), p < 0.01 (**), p < 0.001 (***), p < 0.0001 (****).

Length is a critical property for binding to MHC-I, since the peptide binding groove can usually optimally accommodate only antigenic peptides of specific lengths. Lack of ERAP1 activity has been shown to generally lead to the presentation of longer peptides, consistent with its peptide trimming ability^19,34,38,39^. In both experimental conditions most peptides identified were 9mers, in line with the length preferences of all HLA alleles found in A375 cells (Figure 5B). Still, some notable differences were evident: i) the presence of allotype 10 resulted in the presentation of more 10mers and 11mers and ii) the presence of allotype 2 resulted in the presentation of more 8mers. While both of those observations are consistent with a lower enzymatic activity for allotype 10, comparison with previous studies (Supplemental Figure 2) reveals that the length distribution of the KO cells was significantly more shifted to longer peptides^38,39^, suggesting that allotype 10 retains a significant amount of trimming activity in the cell. In addition to differences in length and predicted affinity, the two sets of peptides differed in the distribution per HLA, with allotype 10 specific peptides being mainly HLA-B*44:03 binders, while allotype 2 specific peptides were more often predicted bind to HLA-A alleles (Figure 5C). These results suggested that ERAP1 allotypic variation can affect both peptide length and HLA prioritization in the cell.

Given the differences between HLA alleles, we further explored this phenomenon by analyzing the predicted affinity and length distribution per HLA (Figure 6). While the tendency for larger peptides for allotype 10 appears to apply for HLA-A and HLA-B alleles (Figure 6D, E,F, J), the smaller predicted affinity for allotype 2 was only significant for A*01:01, A*02:01, B*44:03, C*06:02 and C*16:01 (Figure 6B,C,H,I). Interestingly, allotype 2 generated more 8mers and fewer 9mers than allotype 10, only for HLA-C alleles (Figure 6K-L). Peptides predicted to bind HLA-C alleles also showed the largest difference in affinity between ERAP1 allotypes (Figure 6H-I), including HLA-C*06:02, which has been previously associated with T cell responses in psoriasis^13,48^.

**Figure 6:**
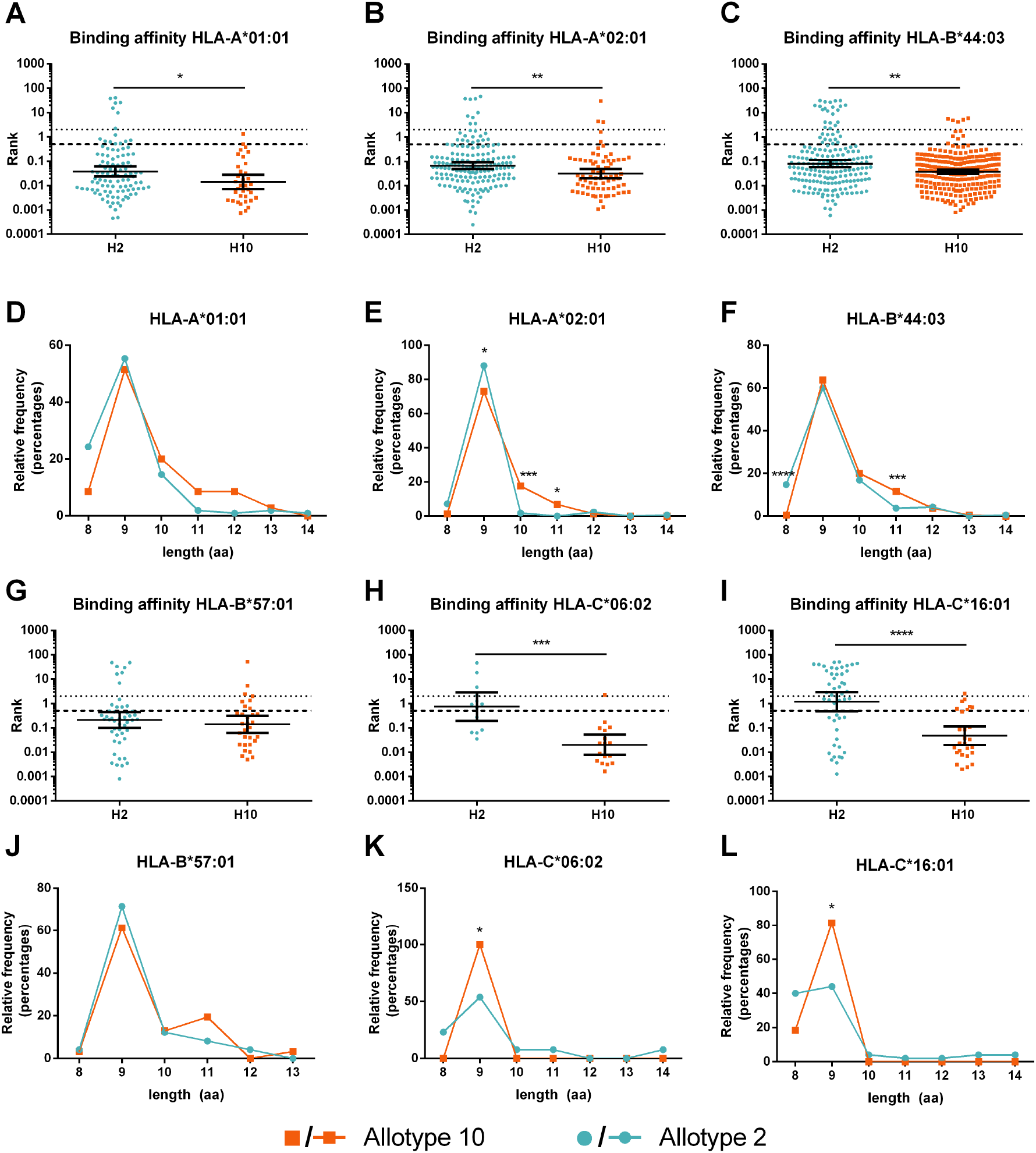
Predicted affinity and length distribution per HLA allele for identified peptides. Each identified peptide was assigned to one of the HLA alleles carried by the cells based on binding predictions. **Panels A-C and G-I**, comparison in predicted affinity for each HLA allele. Statistical significance was evaluated with a Mann-Whitney test in GraphPad Prism v8. p < 0.05 (*), p <0.01 (**), p < 0.001 (***), p < 0.0001 (****). **Panels D-F and J-L**, length distributions of peptides per HLA allele. Statistical significance was evaluated with Fisher’s exact (counts <5) or proportion test with Bonferroni correction for multiple testing. p < 0.05 (*), p < 0.01 (**), p < 0.001 (***), p < 0.0001 (****).

### Differences in sequence motifs

Changes in the sequences of presented peptides between cells expressing different ERAP1 allotypes can result in changes in immunogenicity. To explore this, we compared the sequence motifs per HLA allele for the peptides that were unique or upregulated for each ERAP1 allotype (Figure 7A). While, as expected, most motifs are dominated by the anchor residues that are recognized by each HLA allele, some differences in non-anchor residue positions were evident. Specifically, position 4 for A*02:01 showed a preference for amino acids with negatively charged side chains (D and E) for allotype 10, which was statistically significant for E (Figure 7B). This was also evident, albeit less intense for C*06:02. In addition, and for several alleles, positions 7 and 8 showed a preference for positively charged amino acids (R and K) for allotype 2 (Figure 7A and C-E). These observations suggest that differences in function between ERAP1 allotypes may shape the nature of non-anchor residues of presented peptides. These non-anchor residues are often key for interactions with the T-cell receptor^49^ and inhibitory NK receptors (KIRs)^50^ and can determine the antigenicity of the presented peptides.

**Figure 7:**
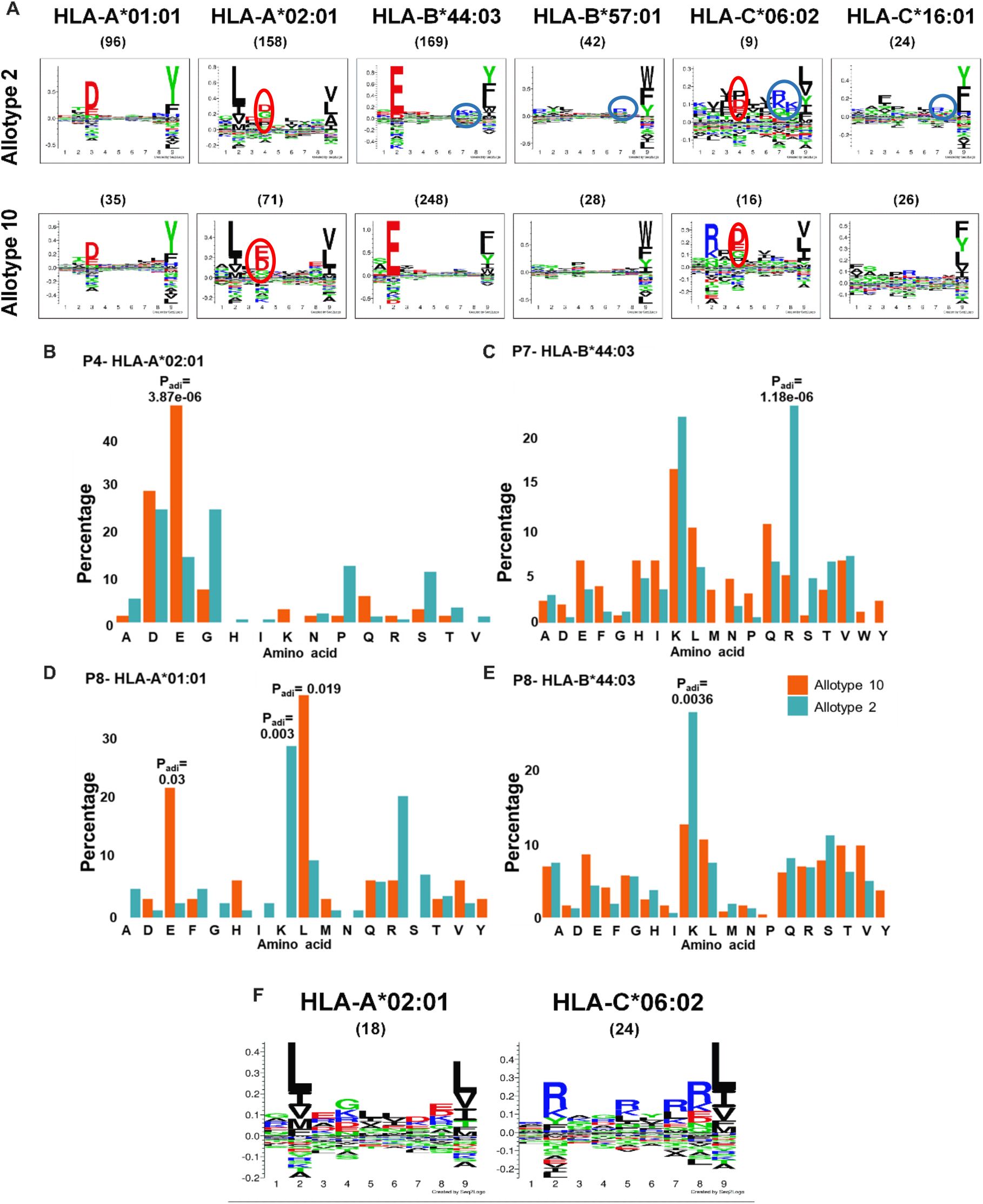
**Panel A**, sequence motifs per HLA allele for peptides unique or upregulated for each ERAP1 allotype. The number of peptides per motif is indicated in parenthesis. Observed differences in motifs are circled. **Panels B-E**, Comparison of amino acid distributions between allotype 2 and allotype 10 cells in P4 for HLA-A*02:01 **(Panel B)**, P7 for HLA-B*44:03 (**Panel C)** and P8 for HLA-A*01:01 **(Panel D)** and HLA-B*44:03 **(Panel E)**. Statistical significance was evaluated with Fisher’s exact (counts <5) or proportion test with Bonferroni correction for multiple testing. **Panel F**, sequence motifs of suspected psoriasis autoantigens from the IEDB database.

Apart from the autoantigenic ADAMTSL5 epitope^13^, several other suspected psoriasis autoantigens exist but were not discovered in our immunopeptidomic analysis. Nevertheless, to examine the potential role of ERAP1 allotypes in the generation of autoantigenic motifs, we downloaded suspected psoriasis autoantigens from the Immune Epitope Database (IEDB)^51^ and used MHCMotifDecon^47^ to extract sequence motifs. Most of the 63 peptide sequences tested were predicted to bind to HLA-A*02:01 and HLA-C*06:02, and therefore we generated sequence logos for these alleles (Figure 7F). This was consistent with the well-established association of HLA-C*06:02 with psoriasis and the reported link between HLA-A*02:01 and psoriasis in Caucasians^52^. In the motifs derived from the suspected psoriasis autoantigens, no D/E amino acid at position 4 was prominent. On the contrary, R/K amino acids at position 7 were common in the HLA-C*06:02 allele. These results suggest that negatively charged residues at position 4 may be less immunogenic in psoriasis, whereas the presence of positively charged residues at positions 7 and 8 may enhance immunogenicity. These findings have very good correlation with the peptide motifs upregulated by allotype 10 and allotype 2, respectively, suggesting that differences in psoriasis predisposition depending on ERAP1 allotypic variation are driven by the generation and presentation of sets of peptides with different immunogenicity.

Immunogenicity towards melanoma cells in cancer is often attributed to the presentation of a set of melanoma-associated antigens (MAGE). To examine if ERAP1 allotypes can affect the presentation of MAGE, we compared the abundance of known MAGE across the two A375 clones (Figure 8). We identified 10 antigenic peptides to be upregulated by allotype 2 and 4 antigenic peptides to be upregulated by allotype 10. Interestingly, some of the peptides were markedly upregulated by allotype 2, such as MAGE A10 which was upregulated by 10-fold. This finding further supports the notion that the two ERAP1 allotypes may induce a distinct immunogenic profile.

**Figure 8:**
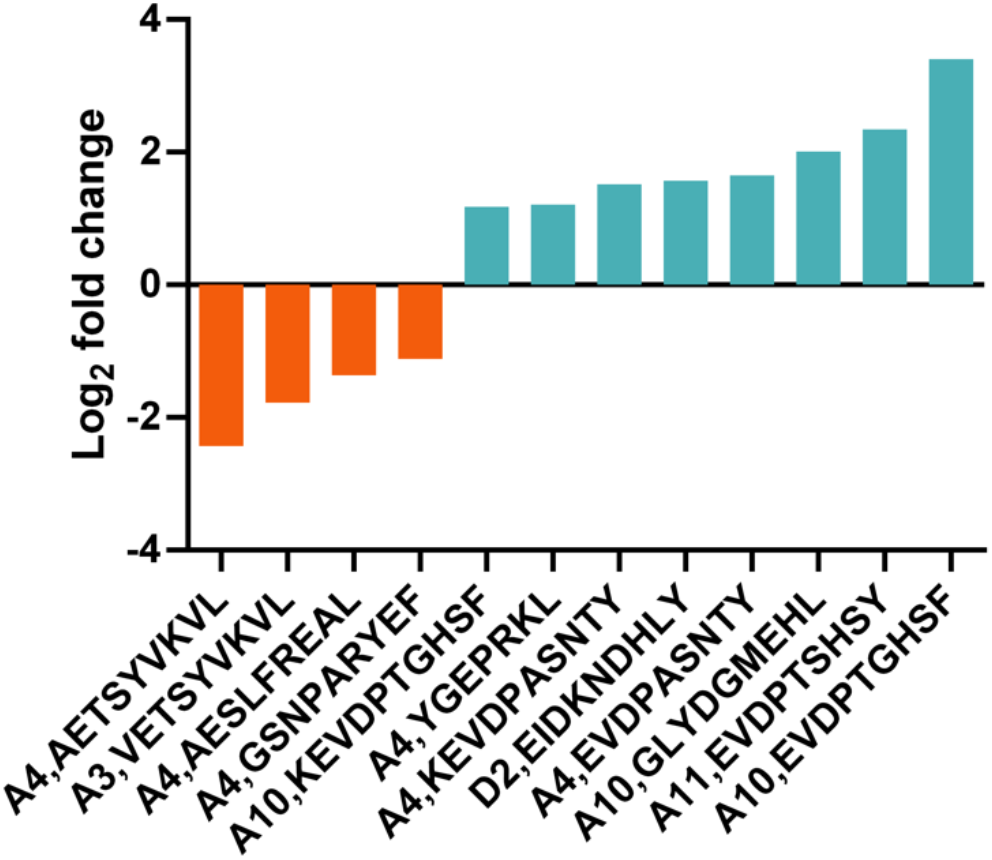
Effect of allotype 2 on the presentation of MAGE-derived antigenic peptides. Log_2_ fold changes in peptide abundance for identified MAGE-derived antigenic peptides are shown. Each peptide sequence and MAGE antigen is indicated.

## DISCUSSION

Several studies have established ERAP1 activity as a key regulator of the cellular immunopeptidome^53,54^, although the extent of the regulation appears to vary among the cellular systems examined. Functional ERAP1 SNPs have also been associated with immunopeptidome shifts, although comparisons between different cell lines have sometimes complicated interpretation^35,55^. These SNPs mainly exist as specific allotypes in the human population and their functional effects often synergize so that allotypes present significantly distinct enzymatic functions. This variability has been associated both with cancer and autoimmunity. Therefore, examining the effect of allotypic variability on immunopeptidome generation is of major importance^30,31,33^.

Among the most common ERAP1 allotypes, allotype 10, the most common in Europeans^30^, is of particular interest, as it was recently demonstrated to downregulate autoreactive immune responses in psoriasis. This was attributed to the inefficient trimming of the ADAMTSL5 antigen, which was however efficiently produced by allotype 2^13,48^. At the same time, in the context of another MHC-I-opathy called Behçet’s disease, allotype 10 enhances autoimmune responses, potentially due to the generation of HLA-B51 restricted peptides, that alter the immunodominance of subsequent CD8+ T cell responses^34^. Although initial functional assays marked allotype 10 as a loss-of-function variant^32^, follow-up *in vitro* analysis by our group suggested more complex changes in activity^31,33^, necessitating deeper analyses.

Our findings here demonstrate a distinct effect by allotypes 2 and 10 on the immunopeptidome that is not consistent with the notion that allotype 10 is a loss-of-function ERAP1 variant. The generation of optimal length 9mers is virtually identical for both allotypes and most changes focus either on the presentation of more 8mers by allotype 2 or more 10mers and 11mers by allotype 10. In addition, the shift in uniquely presented peptides is quite small, since 92% of the detected peptides are common between the two allotypes, a pattern quite distinct from previous comparisons with inhibited or knocked-out ERAP1^19,38^. Rather, the differences focus on relative abundance of identified peptides, in which case the immunopeptidome shift affects 36% of the peptides. These findings are more consistent with two enzyme variants that differ in potency and specificity rather than comparing an active and an inactive variant.

Generation of shorter peptides and destruction of strong binders by over-trimming, observed in allotype 2, may enhance immunogenicity by promoting the dissociation of weaker binding peptides from the cell surface^56^, thereby either allowing the capture of un-edited extracellular peptides or triggering the activation of Natural Killer cells^21^. In parallel, allotype 2 carrying cells were more effective in presenting immunogenic MAGE peptides, some up to 10-fold over allotype 10. Given that antigenic peptide abundance may be an important factor in determining adaptive immune responses^57^, the findings of this study highlight the potential of ERAP1 allotypic variability to drive differences in immunogenicity. Moreover, the extensive changes in the presented peptide repertoire shift the focus from specific antigens to broader immunopeptidome changes.

While the impact of these broad immunopeptidome shifts on adaptive immune responses need to be addressed in future studies, they could be highly relevant in the context of established phenomena such as antigenic drift and T-cell receptor polyspecificity^58,59^. In addition, broad immunopeptidome shifts may affect Natural Killer cell activity, as these cells show a much broader specificity than T cells^50^. Indeed, an HLA tetramer carrying psoriasis-associated antigenic peptides was recently reported to stain CD56+ natural killer cells or KIR2DL1/2DS1 receptors, providing initial evidence for the involvement of NK cells in this autoimmune condition^59^. Interestingly, the side chains at positions 7 and 8 of the presented peptides, and in particular residues R or K, can enhance NK cell activation. This pattern is similar to the allotype 2 motifs detected in our study. Moreover, ERAP1 has been reported to regulate NK cell responses in cancer^21,36,37^ and to specifically regulate KIR receptors engaging HLA-B alleles^60,61^ – which we found to be more extensively utilized by the more immunogenic allotype 2.

While our study defines a framework for understanding the implication of ERAP1 allotypic variation in autoimmunity and anti-tumor responses, it carries some important limitations that should be weighted when interpreting results. Although this study was in part inspired by findings on ERAP1-dependent processing of the ADAMTSL5 antigen in psoriasis, the lack of detection of this peptide and the identification of a low number of HLA-C*06:02 binders, even in the presence of IFN-γ, limits our ability to correlate our findings to that particular autoimmune condition. While this may be an inherent limitation of the A375 cell line, our findings can still contribute to our understanding of HLA-associated autoimmunity and cancer. Further studies utilizing cellular models with different HLA combinations may be necessary to clarify the roles of specific ERAP1 allotypes in different autoimmune conditions. Overall, our results demonstrate a good level of translation of *in vitro* enzymatic properties of ERAP1 to the cellular immunopeptidome, despite the dominant motif-filtering by HLA alleles. Future studies aiming to test the immune effects of these immunopeptidome shifts, including the role of ERAP1 allotypes in disease-specific cells, will be necessary to establish the importance of the observations shown here on the pathogenesis of HLA-associated autoimmunity as well as on anti-tumor immunity.

## FUNDING

Funding was provided by the European Commission in the context of the Marie Skłodowska-Curie Action European Training Network CAPSTONE (954992 – CAPSTONE – H2020-MSCA-ITN-2020). We also acknowledge support of this work by the project “The Greek Research Infrastructure for Personalized Medicine (pMedGR)” (MIS 5002802) which is implemented under the Action “Reinforcement of the Research and Innovation Infrastructure”, funded by the Operational Program “Competitiveness, Entrepreneurship and Innovation” (NSRF 2014-2020) and co-financed by Greece and the European Union (European Regional Development Fund).

## CONFLICT OF INTEREST

All authors declare no commercial or financial conflict of interest.

## AUTHOR CONTRIBUTIONS

**Martha Nikopaschou:** Conceptualization, Methodology, Investigation, Data curation, Formal analysis, Visualization, Writing – review & editing, Writing – original draft. **Martina Samiotaki:** Methodology, Investigation, Data curation (immunopeptidomics). **Anna Kannavou:** Investigation (monoclonal antibody production). **Nikos Angelis:** Methodology, Investigation (flow cytometry). **Ourania Tsitsilonis:** Supervision, Resources (flow cytometry). **George Panayotou:** Resources, Methodology (immunopeptidomics). **Efstratios Stratikos:** Conceptualization, Methodology, Formal analysis, Supervision, Resources, Project administration, Writing – original draft, Writing – review & editing, Funding acquisition.

## Notes

### Competing Interest Statement

The authors have declared no competing interest.

