## Supplemental Figure for "ERAP1 allotypes 2 and 10 differentially regulate the immunopeptidome of melanocytes"

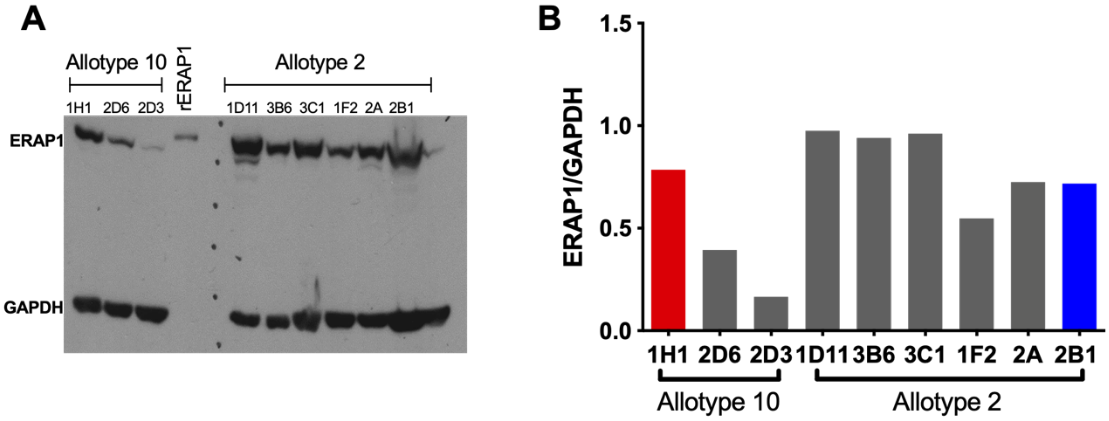


**Supplemental Figure 1: Panel A,** western blot analysis of the obtained clones for ERAP1 expression. rERAP1, recombinant ERAP1 control. GAPDH was used as an internal standard. **Panel B,** densitometric analysis of western blot shown in panel A, showing the ratio of signal for ERAP1/GAPDH as an measure of ERAP1 expression in each clone. Selected clones for further analysis are indicated by red and blue colors.

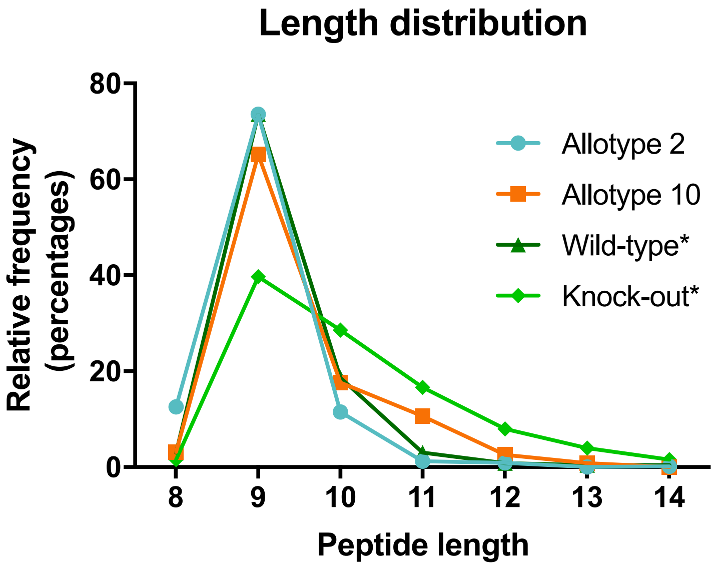


**Supplemental Figure 2:** *Length distribution of identified peptides for the two allotypes in comparison to the distribution for wild-type and ERAP1 KO A375 cells as measured in a previous study (Nikopaschou et al. MCP 2025).*
